# No evidence that synonymous mutations in yeast genes are mostly deleterious

**DOI:** 10.1101/2022.07.14.500130

**Authors:** Leonid Kruglyak, Andreas Beyer, Joshua S. Bloom, Jan Grossbach, Tami D. Lieberman, Christopher P. Mancuso, Matthew S. Rich, Gavin Sherlock, Erik van Nimwegen, Craig D. Kaplan

## Abstract

In a recent paper^1^, Shen *et al*. reported that most mutations in the coding regions of 21 yeast genes were strongly deleterious, and that the distributions of fitness effects were similar for synonymous and nonsynonymous mutations. Taken at face value, these results would conflict with well-established findings from a broad range of fields and approaches. Here, we argue that these results arose from a lack of appropriate controls for the impacts of background genetic effects in edited strains. A re-examination of the data in Shen *et al*. strongly suggests that it is entirely consistent with the expectation that most nonsynonymous and nearly all synonymous mutations have no detectable effects on fitness. We present analyses which show that the data is inconsistent with the proposed explanation that pervasive fitness effects of synonymous mutations arise from their effects on mRNA levels, that the sequence-based fitness assay overestimates fitness effects compared to the growth-based fitness assay, and that the observed wide fitness distributions for nonsense mutations are consistent with ‘off-target’ effects or other uncontrolled sources of biological variation contributing to measured fitness. We conclude by discussing the essential controls and other experimental design considerations that are required to produce interpretable results regarding the fitness effects of mutations in large-scale screens.

## Introduction

Some mutations within a gene’s coding region change the amino acid sequence of the encoded protein; these are known as nonsynonymous mutations. Other mutations alter a gene’s DNA sequence without any changes in the encoded protein; these are known as synonymous mutations. Many lines of evidence accumulated over decades of research have shown that, because they do not lead to sequence changes in proteins, synonymous mutations are much less likely than nonsynonymous ones to have biological effects. However, in a recent paper^1^, Shen *et al*. reported that most mutations in the coding regions of 21 yeast genes were strongly deleterious, and that the distributions of fitness effects were similar for synonymous and nonsynonymous mutations. Taken at face value, these results would conflict with well-established findings from a broad range of fields and approaches, including human genetics^2–7^, population genetics^8–10^, mutagenesis screens^11–13^, and deep mutational scanning^14–16^. These prior findings have firmly established that only some nonsynonymous changes have detectable fitness effects, and that synonymous changes are much more likely to be neutral than nonsynonymous ones. These prior findings are highly robust, despite evidence that some synonymous changes can have detectable fitness consequences because they create or disrupt functional elements embedded within coding sequences^17^ or allow for tuning of translational properties and feedback on mRNA stability^18–20^. For additional context and analysis, please see the accompanying letter^21^.

Here, we argue that there is an alternative explanation for the data presented by Shen *et al*. Flaws in experimental design and analysis—most importantly a lack of appropriate controls for the impact of background genetic effects in edited strains—make their interpretation that the majority of both synonymous and nonsynonymous mutations are strongly deleterious untenable. Instead, we predict that inclusion of appropriate controls would show that the results are completely consistent with the expectation that most nonsynonymous and nearly all synonymous mutations have no or minimal effects on fitness; only a subset of mutations, primarily nonsynonymous ones, would have significant effects on fitness. We discuss what controls are needed to be able to properly assess the fitness effects of mutations in large-scale CRISPR editing screens.

## Results

Shen *et al*. set out to examine fitness effects of synonymous and nonsynonymous mutations in a curated set of 21 genes in yeast^1^. They first used CRISPR-Cas9 to create 21 strains each carrying a deletion of 150 nucleotides and a common sgRNA target in one of the genes. They then used CRISPR-Cas9 to generate a double-stranded break at the deletion site, which was repaired using pools of synthesized templates homologous to sequences flanking each deletion (Illustrated in Figure 1). This process would hypothetically generate libraries of all 450 single-nucleotide variants in the chosen 150 nucleotide region of each gene, although in the published experiment, some variants were missing or lacked sufficient DNA sequencing coverage for analysis. Shen *et al*. then subjected these libraries to competitive growth, with a single “wildtype” (WT) strain, derived from repair of a strain carrying a deletion in one of the genes (*ASC1*), mixed into the libraries as a control. The fitness of each strain was determined from its relative abundance over time compared to this WT strain, measured by deep sequencing of amplicons for the mutated regions of each gene. Most mutant strains showed a relative depletion over time, leading to the reported results. The key results of Shen et al. are presented in their Figure 2c, which appears to show that for each gene, the distribution of fitness effects was similar for synonymous and nonsynonymous edits. We note that for about half of the genes, a subset of edits shows much stronger deleterious effects, and that nearly all of these edits are nonsynonymous.

**Figure 1.**
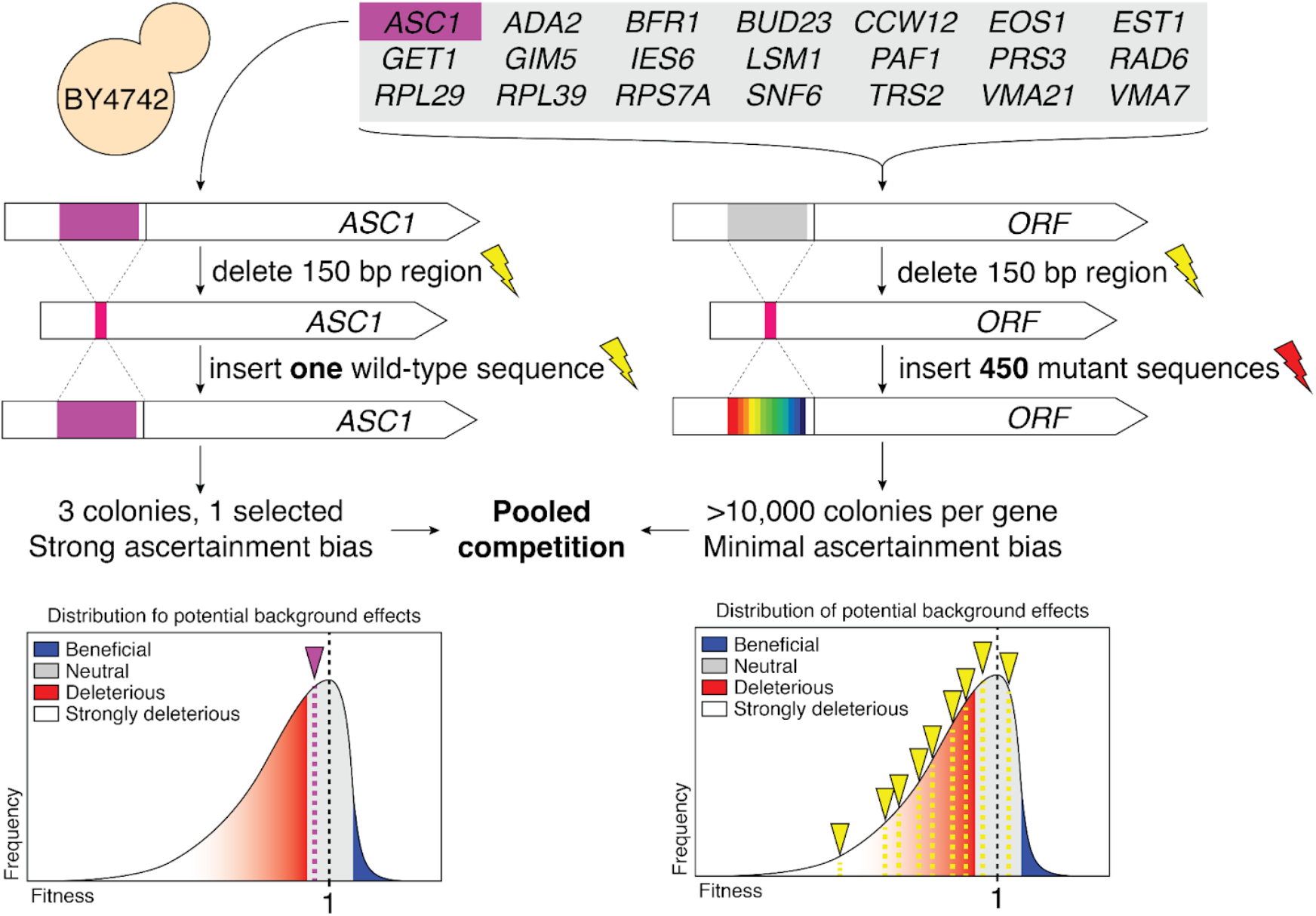
Schematic of the approach taken by Shen *et al*. for strain creation. Parallel strain construction allows for background effects on fitness (indicated by lightning bolts) that may be different between strains (yellow bolt vs red bolt). Furthermore, selection of a single wild-type strain (left) creates uncontrolled ascertainment bias relative to variant strains that will incorporate the average of background effects across a distribution (right) into any fitness measurement.

**Figure 2.**
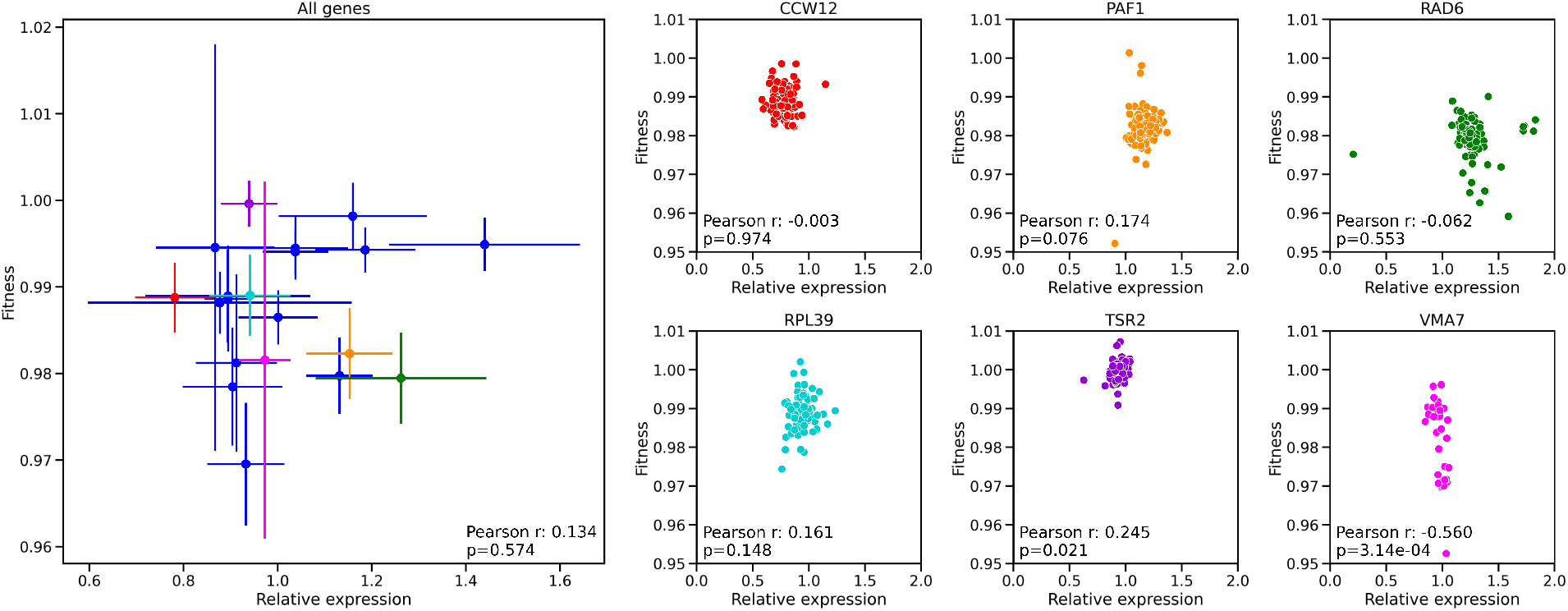
(Left) Means and standard deviations of the fitness (vertical axis) and relative mRNA levels of synonymous edits (horizontal axis) across genes. Although the distributions of the effects on fitness and relative mRNA levels of synonymous edits vary substantially across genes, there is no correlation between the average effect on fitness and the average effect on relative mRNA level (Pearson correlation coefficient 0.134 with *p*-value 0.57 under a Pearson correlation test; see Methods for description of error calculation). (Right) There is little correlation between the fitness and relative expression of synonymous mutations within genes. A subset of genes is shown; most synonymous mutations cluster around the mean fitness and relative expression of the gene. Points are mean fitness and expression values for all variants with 4 replicates of each measurement.

In addition to the fact that the reported results contradict a large body of prior work, we note a number of inconsistencies in the interpretation offered by Shen *et al*. First, variants with deleterious effects as large as those reported in Figure 2a-c should be efficiently removed by purifying selection; however, Figure 2e shows that many of them are observed in other yeast species. Second, Shen *et al*. propose that the pervasive deleterious effects of synonymous mutations arise from their impact on gene expression levels. However, while the distributions of the effects of synonymous mutations on fitness and on mRNA levels vary substantially across genes (see Shen *et al*. Fig. 2c and Extended Data Fig. 4), there is no correlation between the average effects on fitness and on mRNA levels (Figure 2). For some genes *(e.g., VMA7),* virtually all synonymous mutations are reported to be strongly deleterious, but their effects on mRNA levels are very small. These inconsistencies are easily explained by our alternative interpretation of the data, *i.e.,* that most of the reported fitness effects arise not from the mutations in question, but from other effects of the genomic background of each strain, which we discuss below.

Crucially, the repair template pools used by Shen *et al*. did not contain any WT sequences that would allow for WT versions of each gene to be created in a manner identical to and in parallel with the creation of the genome-edited strains. Inclusion of such WT strains in the libraries would have allowed for proper control of any background effects on fitness that are specific *for each* edited gene.

Several critical factors make such controls essential for determining the fitness of each edited strain. First, transformation of yeast itself can be mutagenic^22,23^, and making deletions in each gene with CRISPR-Cas9 can generate off-target effects, with different guide RNAs causing different types of off-target effects and with different frequencies. These experimental manipulations mean that each genetically engineered deletion strain represents a unique genetic background potentially carrying unascertained mutations that may affect strain fitness (Figure 1).

Second, the effects of deleting particular genes may have caused other mutations to emerge during growth, before rescue with the variant library. For example, deletion of *RAD6* is expected to impair DNA repair and may lead to chromosomal rearrangements^24^, deletion of *EST1* is expected to alter telomere length and function^25,26^, and deletion of *PAF1* is expected to cause defects in multiple histone modifications^27,28^. Therefore, it is not clear that each of these deletions, after some period of growth, would necessarily be complemented immediately to WT fitness upon reintroduction of a WT allele, let alone upon introduction of a mutant allele. The combination of all of these effects means that strains in which each of the 21 genes were first deleted may have fitness differences from each other and from the single WT control strain, making it impossible to properly assign fitness to the individual engineered mutations. Differences between the 21 genetic backgrounds could have been examined with wholegenome sequencing of the strains, but such sequencing was not carried out.

Third, further unascertained fitness-altering mutations may have arisen during CRISPR-Cas9-mediated insertion of variant alleles, contributing to the observed fitness differences among different edits of each gene. Critically, in pooled library construction, each individual variant derives from many colonies, and these colonies sample a distribution of fitness effects arising from any of the above-mentioned variant-independent mechanisms; the reported fitness for each variant then reflects the average of this distribution (Figure 1). By contrast, the individual WT colony selected to serve as the control does not represent an average of the distribution of potential fitness effects from these allele-independent mechanisms, and indeed may have been selected as a robustly growing colony, which is much more likely to have wild-type fitness.

Fourth, careful reexamination of the reported growth rates for reconstructed synonymous variants in Figure 1d of Shen *et al*. (Figure 3) shows that relative growth rates of most variants cluster around 1, while fitness determination from pooled growth competition indicated that fitness of these variants was mostly below 1 (slope of 0.89 for the linear fit). These results are indicative of potential bias in one of the assays.

**Figure 3.**
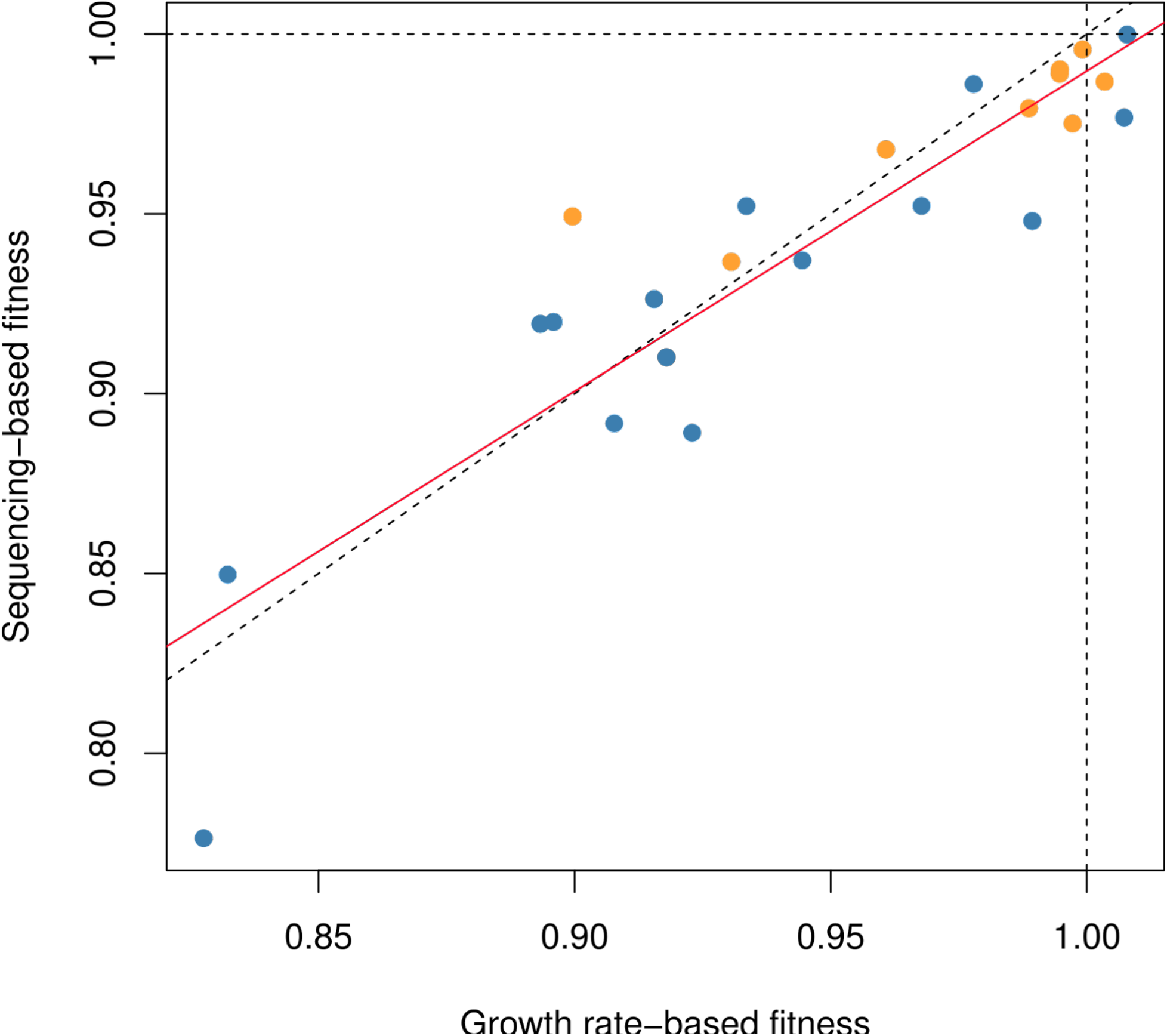
Comparison of sequencing-based fitness and growth rate-based fitness for synonymous variants (orange) and non-synonymous (blue). The deviation from a slope of 1 is consistent with an underestimation of fitness in the sequence-based fitness assay, and therefore an overestimation of small effects, compared to the growth-based fitness assay (dashed line represents x=y, red line is linear fit, slope=0.89).

Diagnosis and alleviation of these technical issues could have been accomplished through the use of many independent isolates for each of the 21 deletions, and through the generation of multiple WT alleles in each of the deletion backgrounds in biological replicates, but such biological replicates were not performed. We propose that the observed differences in average fitness among the 21 genes in Figure 2c of Shen *et al*. represent such uncontrolled effects of genetic background. The selection of a single WT strain is especially sensitive to potential ascertainment bias, as any obviously unhealthy strain would not be selected to represent WT fitness, but within the variant pools and among pools of the same variant there is no such filter.

Indeed, the results shown in Shen *et al*. Figure 2c are completely consistent with such artifacts. Most of the genes show an overall average impaired fitness relative to the single WT control, and these values differ across the 21 genes. For each gene, the mean and the distribution of fitness are very similar for synonymous and nonsynonymous edits; we expect that they would also be similar for control edits that simply re-introduced the wild-type sequences for each gene, had such appropriate gene-specific controls been used. Roughly half of the genes have a small subset of edits with much larger fitness effects that clearly stand out from the rest of the distribution, and nearly all of these edits are nonsynonymous (see Figure 4 for fitness distributions of six example genes). We expect that these represent the real deleterious effects of these nonsynonymous mutations, with the reported fitness values of the rest of the edits for each gene inadvertently serving as the true null distribution. We also found that nonsense mutations present in the experiment often exhibit a wide distribution of fitness effects (Fig. 4, blue curves), further supporting our interpretation that off-target effects make major contributions to measured fitness. We would expect that true fitness of effects of different nonsense mutations in a given gene should generally be equivalent notwithstanding potential partial loss of function due to tolerated truncations. Consistently, the magnitude of observed fitness effects was not correlated with the position of nonsense mutations within the edited regions.

**Figure 4:**
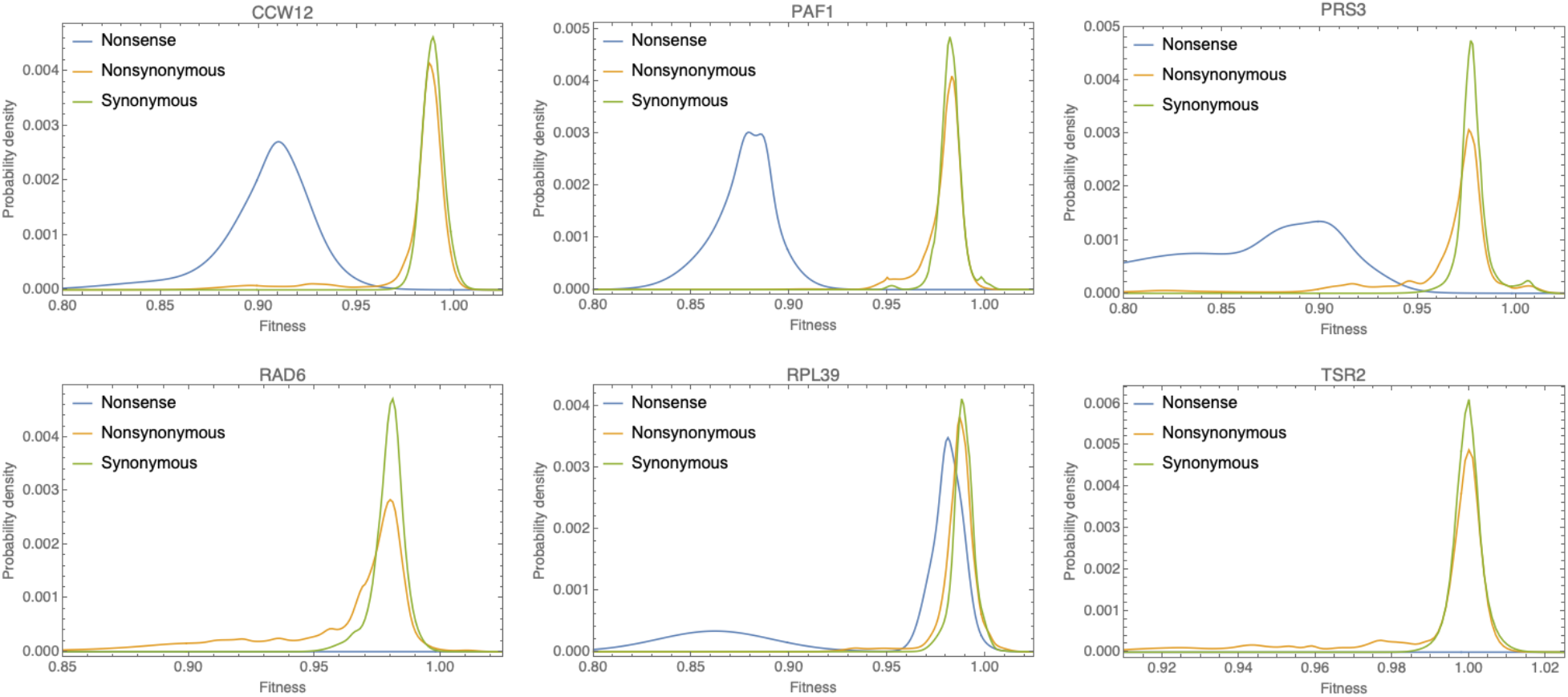
Distributions of measured fitness for synonymous (green), nonsynonymous (orange) and nonsense (blue) edits for six example genes (see panel headers) that exhibit a tail of nonsynonymous edits with strong deleterious effects. Note that different nonsense edits of the same gene often exhibit substantially different fitness, leading to wide fitness distributions of nonsense mutations. These distributions do not correlate with locations of nonsense alleles within the edited regions. This observation is consistent with ‘off-target’ effects or biological variation other than that of the nonsense mutation contributing to the measured fitness. Note that no nonsense edits were measured for *RAD6* and *TSR2*.

## Conclusions

To settle these issues empirically, the experiment requires repetition with the inclusion of WT sequences in each variant pool. Without this indispensable control, the results are largely uninterpretable. Should the experiment be repeated with the use of such control sequences, the following important points need to be incorporated to make any conclusions as robust as possible. Growth measurements for multiple independent isolates of each deletion strain are needed to allow determination of potential off-target effects and deletion-specific fitness. Wholegenome sequencing of each isolated deletion strain should be carried out in order to uncover potential cryptic alleles created during strain construction. In addition, biological replicates should be employed during the creation of variant libraries by genome editing to reliably measure the fitness effect, if any, for each variant; otherwise, these cannot be distinguished from off-target effects during library construction. In the configuration employed by Shen *et al.,* fitness measurements from replicate competitions of the same pool represent technical replication of competition only. As a confirmatory step, direct fitness measurements of many independently created individual variants of a gene in parallel with WT alleles created on the identical genetic background should be performed. More generally, we note that when results appear to contradict decades of evidence from multiple fields, the default assumption should be that they arise from a hidden artifact or artifacts, and every effort should be made to rule these out before taking the results at face value.

## Methods

### Fitness distributions

Fitness data were taken from Shen *et al*. Supplemental Data 3^1^. For each gene, edits were stratified into three classes: synonymous, non-synonymous, and nonsense. Only mutations with at least two replicate measurements were retained. For each edit the mean fitness and standard-deviation were calculated from the replicate measurements and the fitness distribution for each class was estimated as the mixture of normal distributions with corresponding means and standard-deviations for all edits within the class (shown in Figure 4).

### Comparison of the average effects of synonymous mutations on fitness and mRNA levels

Relative expression data were taken from Shen *et al*. Supplemental Data 4^1^. For each gene, all synonymous edits with at least two replicate measurements were taken, and the mean and standard deviation across the replicates were calculated for each edit. As above, we estimated the overall fitness distribution of synonymous mutations as the mixture of normal distributions with the corresponding means and standard deviations and the overall mean and standard deviation of this mixture were determined for each gene. For the mRNA levels an overall mean and standard deviation was calculated analogously for each gene. The means and error-bars in Figure 2 correspond to these means and standard-deviations of the effects on fitness and mRNA level of synonymous edits for each gene. For gene-specific correlations in Figure 2 (Right), datasets were filtered for synonymous variants that were assayed in all four replicates for both fitness and relative expression. Pearson correlations and p-values were calculated using the *pearsonr* function from Scipy^29^.

### Comparison of fitness assays

Fitness measurements based on growth- and sequencing-based assays were taken from the supplemental data of Shen *et al*. (Source Data Fig.1)^1^. To quantify the linear relationship between the estimates generated by the two assays, as shown in Figure 3, we built a linear model using the *lm*-Function of the R-stats package^30^. We modeled sequencing-based fitness as the dependent variable y with a freely chosen offset and the growth-based fitness as the only predictor *x:*

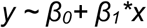

